# ASIC3-dependent spinal cord nociceptive signaling in cutaneous pain induced by lysophosphatidyl-choline

**DOI:** 10.1101/2020.12.28.424561

**Authors:** Ludivine Pidoux, Kevin Delanoe, Eric Lingueglia, Emmanuel Deval

**Affiliations:** Université Côte d’Azur, CNRS, IPMC, LabEx ICST, FHU InovPain, France

## Abstract

Lysophosphatidyl-choline (LPC), a member of the phospholipid family, has recently emerged as an interesting new player in pain. It has been proposed to mediate pain through Acid-Sensing Ion Channel 3 (ASIC3), a pain-related channel mainly expressed in peripheral sensory neurons. LPC potentiates ASIC3 current evoked by mild acidifications, but can also activate the channel at physiological pH, and its local injection in rodents evokes ASIC3-dependent pain. We combine here *in vivo* recordings of spinal cord neuron activity with subcutaneous LPC injection to analyze the mechanism of action associated with the LPC-induced, ASIC3-dependent pain in peripheral and spinal cord neurons. We show that a single cutaneous injection of LPC exclusively affects the nociceptive pathway. It evokes an ASIC3-dependent short-term sensitization of nociceptive fibers that drives hyperexcitability of projecting neurons within the dorsal spinal cord without apparent central sensitization.

## INTRODUCTION

Lysophosphatidyl-choline (LPC) is an endogenous lysolipid produced following plasma membrane hydrolysis by PLA2 enzymes (Murakami *et al*., 2020) and has recently emerged as an interesting new player in pain (Hung *et al*., 2020; Marra *et al*., 2016; Rimola *et al*., 2020). We already identified high levels of LPC in synovial fluid from patients with painful joint diseases (Marra *et al*., 2016), and increased LPC was recently detected in the plasma of a subset of patients with fibromyalgia (Hung *et al*., 2020). In all these conditions, LPC has been proposed to mediate pain through peripheral Acid-Sensing Ion Channels 3 (ASIC3), which are activated and potentiated by this lysolipid.

Acid-Sensing Ion Channels (ASICs) are a family of cation channels identified in the late 90’s, which are known for their ability to sense extracellular protons (Waldmann *et al*., 1997b). Several ASIC subunits have been cloned in mammals, including ASIC1, ASIC2, ASIC3 and ASIC4, with variants (for reviews, see (Deval *et al*., 2010; Lee and Chen, 2018). A functional ASIC channel results from the trimeric assembly of these subunits (Jasti *et al*., 2007), at least of ASIC1, ASIC2 and ASIC3, leading to homomeric and heteromeric channels with different biophysical properties (Hesselager *et al*., 2004). ASICs are basically considered as extracellular pH sensors but their activity and/or expression are also highly regulated by various endogenous factors associated with ischemia, inflammation and pain (Allen and Attwell, 2002; Birdsong *et al*., 2010; Deval *et al*., 2008; Deval *et al*., 2004; Immke and McCleskey, 2001; Li *et al*., 2010; Mamet *et al*., 2002; Mamet *et al*., 2003; Sherwood and Askwith, 2009; Smith *et al*., 2007; Xiong *et al*., 2004). This is particularly the case for ASIC3 channels (Waldmann *et al*., 1997a), which have been reported to behave as “coincidence detector” of mild extracellular acidifications, hypertonicity, arachidonic acid and/or ATP (Birdsong *et al*., 2010; Deval *et al*., 2008; Li *et al*., 2010). More recently, we showed that LPC, alone or in combination with arachidonic acid (AA), activates a small but non-inactivating ASIC3 current at physiological pH7.4, in addition to its potentiation of the current evoked by mild acidifications (Marra *et al*., 2016).

Peripheral injection of LPC together with AA in rodent skin was shown to generate ASIC3-dependent acute pain behaviors (Marra *et al*., 2016). Understanding how this local injection of LPC affects the nociceptive pathway and the activity of dorsal spinal cord neurons, as well as the contribution of ASIC3 to this process, is therefore of particular interest. We perform here *in vivo* experiments to analyze the mechanism of action associated with cutaneous LPC effect, determine the role of peripheral ASIC3 to the generation of the pain message, and study how this message is integrated at the spinal cord level. We show that LPC, which activates and potentiates ASIC3 *in vitro*, positively modulates both the spontaneous and evoked activities of particular subsets of dorsal spinal cord neurons. Indeed, cutaneous injection of LPC in rats or mice enhances the firing of high threshold (HT) and wide-dynamic range (WDR) neurons, leaving the activity of low threshold (LT) neurons unaffected. Effects of cutaneous LPC injection are associated to short-term sensitization of nociceptive fibers, are significantly reduced by the pharmacological inhibition of ASIC3, and almost completely abolished in ASIC3 knockout mice. This work shows how a single cutaneous administration of LPC provokes peripheral sensitization of ASIC3-expressing nociceptive fibers that drive hyperexcitability of projecting neurons within the dorsal spinal cord.

## RESULTS

### In vivo cutaneous injection of LPC affects spinal nociceptive neurons but not sensory ones

LPC is able to positively modulate ASIC3 (Fig. 1 and Supplementary Fig. 1) by increasing the channel sensitivity to protons, leading to the generation of a constitutive current at resting physiological pH7.4 (Marra *et al*., 2016). It has also been associated to acute pain behaviors when injected locally in rodents (Marra *et al*., 2016). We thus performed *in vivo* recordings of spinal cord dorsal horn neuron activity to study how the acute pain message generated by subcutaneous injection of LPC is integrated at the spinal cord level (Fig. 1A).

**Figure 1:**
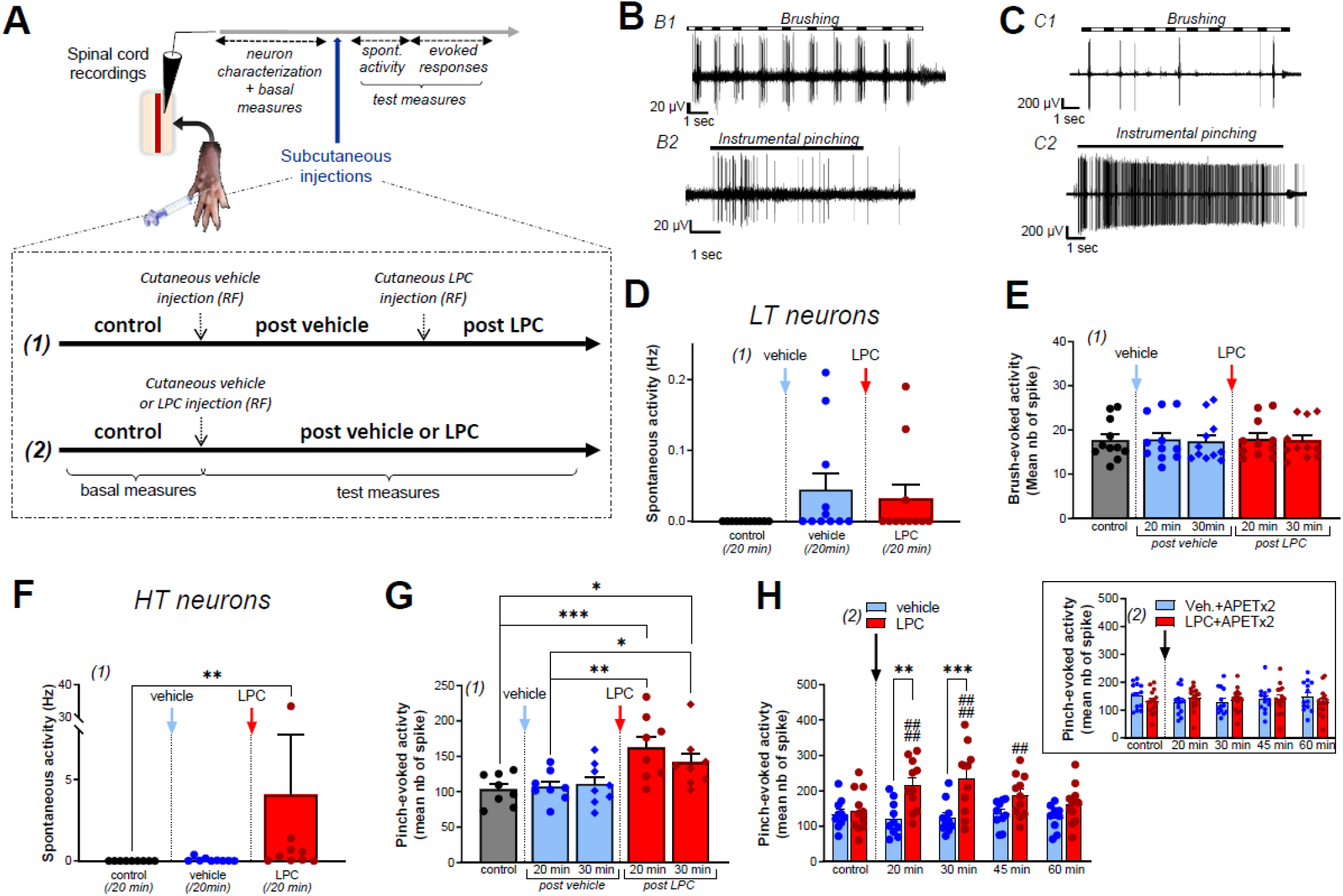
Effect of LPC cutaneous injection on low threshold (LT) and high threshold (HT) neuron activities in rats. **(A)** Protocol used for spinal cord neuron recordings. After neuron characterization in the spinal cord, vehicle and/or LPC were subcutaneously injected in the hindpaw using two protocols described in the inset. **(B)** Typical discharge of a low threshold (LT) neuron responding to non-noxious brushing (B1), but not to noxious pinching (B2, instrumental pinching, 300g). **(C)** Typical discharge of a high threshold (HT) neuron that did not respond to brushing (C1) but emitted a sustained discharge in response to pinch (C2, instrumental pinching, 300g). **(D)** Global spontaneous activity of LT neurons measured with protocol (1) over 20 min-periods following vehicle (blue bar and points) and LPC16:0 (red bar and points) subcutaneous injection within the neuron’s receptive fields (n= 11 neurons from 7 rats, Friedman test with p=0,0988). **(E)** Brushing-evoked response of LT neurons before (control) and after vehicle/LPC16:0 subcutaneous injection in their receptive fields (n= 11 neurons from 7 rats, no significant differences with p=0.9822, Friedman test; protocol (1) used). **(F)** Global spontaneous discharge of HT neurons after vehicle (blue bar and points) and LPC16:0 (red bar and points) injection (n=8 neurons from 7 rats, Friedman test with p<0.0001 followed by a Dunn’s multiple comparison test: **, p=0.0012; protocol (1) used). **(G)** Pinching-evoked responses of HT neurons (instrumental pinching, 300g) before and after vehicle/LPC16:0 subcutaneous injection in their receptive fields (n=8 neurons from 6 rats, Friedman test with p<0.0001 followed by a Dunn’s multiple comparison test: * p<0.05, ** p<0.01 and *** p<0.001; protocol (1) used). **(H)** Duration of the LPC effect on the pinching-evoked activity of HT neurons (n=10 and 11 neurons for vehicle and LPC, respectively; two-way ANOVA test with p=0.0156 and p<0.0001 for treatment and time after injection, respectively; **, p<0.01 and ***, p<0.001, Sidak’s multiple comparison post hoc test; ##, p<0.01 and ####, p<0.0001 as compared to control before LPC injection, Dunnet’s multiple comparison post hoc test; protocol (2) used). The potentiation by cutaneous LPC was still significant 45 minutes after injection. Inset: The effect of LPC was abolished when APETx2 (0.2 nmol) was co-injected together with the lipid. Note that APETx2 had no effect itself (vehicle + APETx2) on the pinching-evoked activity of HT neurons (n=13 and 15 for vehicle+APETx2 and LPC+APETx2, respectively; two-way ANOVA test with p=0.8062 and p=0.7850 for treatment and time after injection, respectively; protocol (2) used).

To determine whether LPC affects the firing of spinal cord neurons, both spontaneous activity and evoked neuronal responses to non-noxious and/or noxious stimulations were monitored in rats before and after vehicle or LPC injections (see Methods, Fig. 1A-C and Fig. 2A). Two different protocols were used for vehicle and LPC delivery (Fig. 1A): (1) consecutive delivery of vehicle and LPC on same animals, which allowed making paired analyses, and (2) alternative delivery of vehicle or LPC on different animals so that they were only injected once. With protocol (1), no significant effect of LPC on low threshold (LT) neurons was observed on either the spontaneous activity (Fig. 1D, 0.04 +/- 0.02 Hz and 0.03 +/-0.02 Hz after vehicle and LPC injections, respectively), or the non-noxious brush-evoked activity (Fig. 1E). The spiking activity evoked by brushing remained unchanged after 20 and 30 minutes of vehicle or LPC injections, as compared to the activity evoked in control condition before injections.

**Figure 2:**
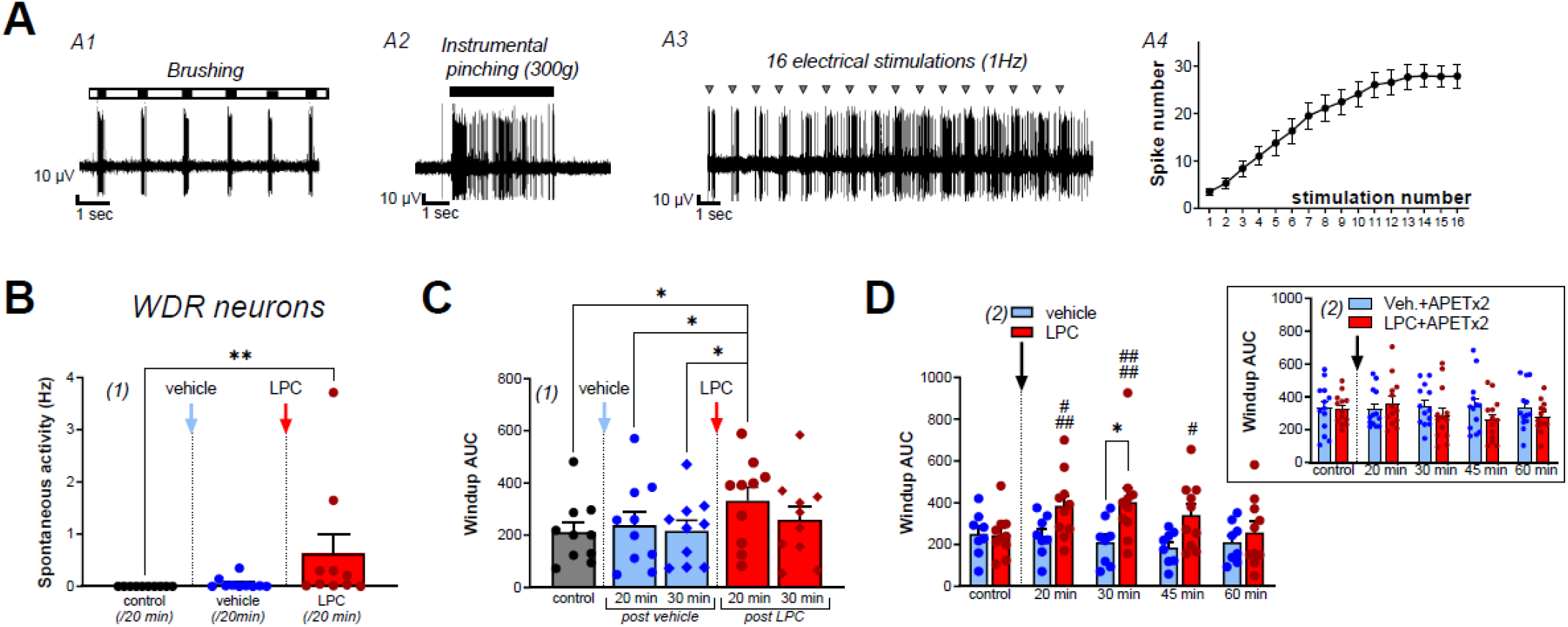
Effect of LPC cutaneous injection on wide dynamic range (WDR) spinal neuron activity in rats. **(A)** Typical discharge of a wide dynamic range (WDR) neuron, which responded to both brushing (A1) and pinching (A2, instrumental pinching, 300g). WDR neuron also exhibited windup (A3) following repetitive electrical stimulations of its receptive field (grey triangle). Typically, windup is characterized by a progressive increase in the number of C-fiber evoked spikes as the number of stimulations increase (A4, data from 22 neurons). Maximal windup was reached between the 13^th^ and 16^th^ stimulation, from 3.48±0.72 spikes at the 1^st^ stimulation to 27.86±2.5 spikes at the 16^th^ stimulation. **(B)** Global spontaneous discharge of WDR neurons after vehicle (blue bar and points) and LPC16:0 (red bar and points) subcutaneous injection in their receptive fields (n=10 neurons from 8 rats, Friedman test with p=0.0008 followed by a Dunn’s multiple comparison test: **, p=0.0036). **(C)** WDR neuron windup before and after vehicle (blue bar and points) and LPC16:0 (red bar and points) subcutaneous injection in their receptive fields. C-fiber induced windup is represented as the area under curve (AUC) determined from classical windup curves (see Methods, n=10 neurons from 8 rats, Friedman test with p=0.0111 followed by a Dunn’s multiple comparison test: *, p<0.05). **(D)** The potentiating effect of LPC on WDR neuron windup was significant for 45 minutes (n=10 neurons from 8 rats and n=8 neurons from 7 rats for LPC and vehicle, respectively, two-way ANOVA test with p=0.0835 and p=0.0047 for treatment and time after injection, respectively; *, p<0.05, Sidak’s multiple comparison post hoc test; ##, p<0.01 as compared to control before LPC injection, Dunnet’s multiple comparison post hoc test), and was abolished when APETx2 (0.2 nmol) was co-injected together with LPC. Note that APETx2 had no effect itself (vehicle + APETx2) on WDR windup (inset: n=13 and 13 for vehicle+APETx2 and LPC+APETx2, respectively; two-way ANOVA test with p=0.4594 and p=0.3816 for treatment and time after injection, respectively).

LPC was next tested on high threshold (HT) (Fig. 1F-H) and wide dynamic range (WDR) (Fig. 2B-D and Fig. 2 and Supplementary Fig. 2) neurons. The spontaneous activities of these 2 neuron types were both significantly increased by LPC cutaneous injection, as compared to vehicle (Fig. 1F, HT neurons: 0.07±0.04 Hz for vehicle *vs*. 4.05±3.67 Hz for LPC; Fig. 2B, WDR neurons: 0.02±0.01 Hz for vehicle *vs*. 0.66±0.41 Hz for LPC). Moreover, the pinch-evoked activity of HT neurons was significantly increased by LPC as compared to control and vehicle conditions (Fig. 1G, +4.5%, and +7.2% compared to control at 20 and 30 minutes, respectively, after vehicle injection, and +57.7% and +36.0% compared to control at 20 and 30 minutes, respectively, after LPC injections). These results were confirmed using protocol (2) in which vehicle or LPC were alternatively delivered to different batches of animals. HT neuron hyperexcitability was induced by LPC and lasted up to 45 minutes after its cutaneous injection, showing short-term sensitization of noxious mechanical sensitivity (Fig. 1H), which was abolished when the ASIC3 blocker APETx2 (Diochot *et al*., 2004) was co-injected together with LPC (Fig. 1H, inset). Similarly, the C-fiber evoked activity of WDR neurons (Supplementary Fig. 2A) was also enhanced by LPC, as illustrated by its effect on windup (Fig. 2C-D and Supplementary Fig. 2A and 2C), whereas no effect was observed on the WDR activities related to Aβ or Aδ fibers (Supplementary Fig. 2A, 2B and 2D). LPC significantly increased windup as compared to vehicle and to control conditions (Fig. 2C, +13.0% and +2.9% compared to control at 20 and 30 minutes, respectively, after vehicle injections, and +57.7% and +22.7% compared to control at 20 and 30 minutes, respectively, after LPC injection), with an effect duration of at least 45 minutes (Fig. 2D). Finally, and as observed for the LPC potentiation of HT neuron evoked-activity, the potentiating effect on WDR neuron windup was also abolished by APETx2 (Fig. 2D, inset).

These results showed that LPC did not impact spinal LT neuron basal and evoked activities in rats, demonstrating that Aβ peripheral fibers were not affected by its cutaneous injection. Conversely, LPC induced hyperexcitability in both HT and WDR spinal neurons that was driven by C-fiber inputs, most probably through an activation of ASIC3 channels as illustrated by the effect of APETx2.

### The LPC-induced mechanical sensitization of spinal nociceptive neurons depends on peripheral ASIC3 activity

Both the spontaneous and evoked activities of mice HT neurons were also significantly potentiated following LPC cutaneous injection (Fig. 3A-B) as observed in rats, whereas those of LT neurons remained unaffected (Supplementary Fig. 3A-B). HT neuron spontaneous activity was significantly higher after LPC injection as compared to vehicle (Fig. 3A, 0.23+/-0.12 Hz for vehicle *vs*. 0.53 +/-0.17 Hz for LPC). Moreover, response of mice HT neurons to noxious pinching was also enhanced by LPC, with a 52.4 % and 53.9% increase in evoked-activity at 20 and 30 minutes after lipid injection, respectively, as compared to vehicle (Fig. 3B). The potentiating effect of LPC lasted up to 45 minutes (Supplementary Fig. 3E), similarly to what was observed in rats. Importantly, all the effects on both the spontaneous and pinch-evoked activities were lost in ASIC3 knockout mice (ASIC3 KO, Fig. 3C-D), demonstrating the involvement of ASIC3 channels in the LPC-induced hyperexcitability of nociceptive HT spinal neurons.

**Figure 3:**
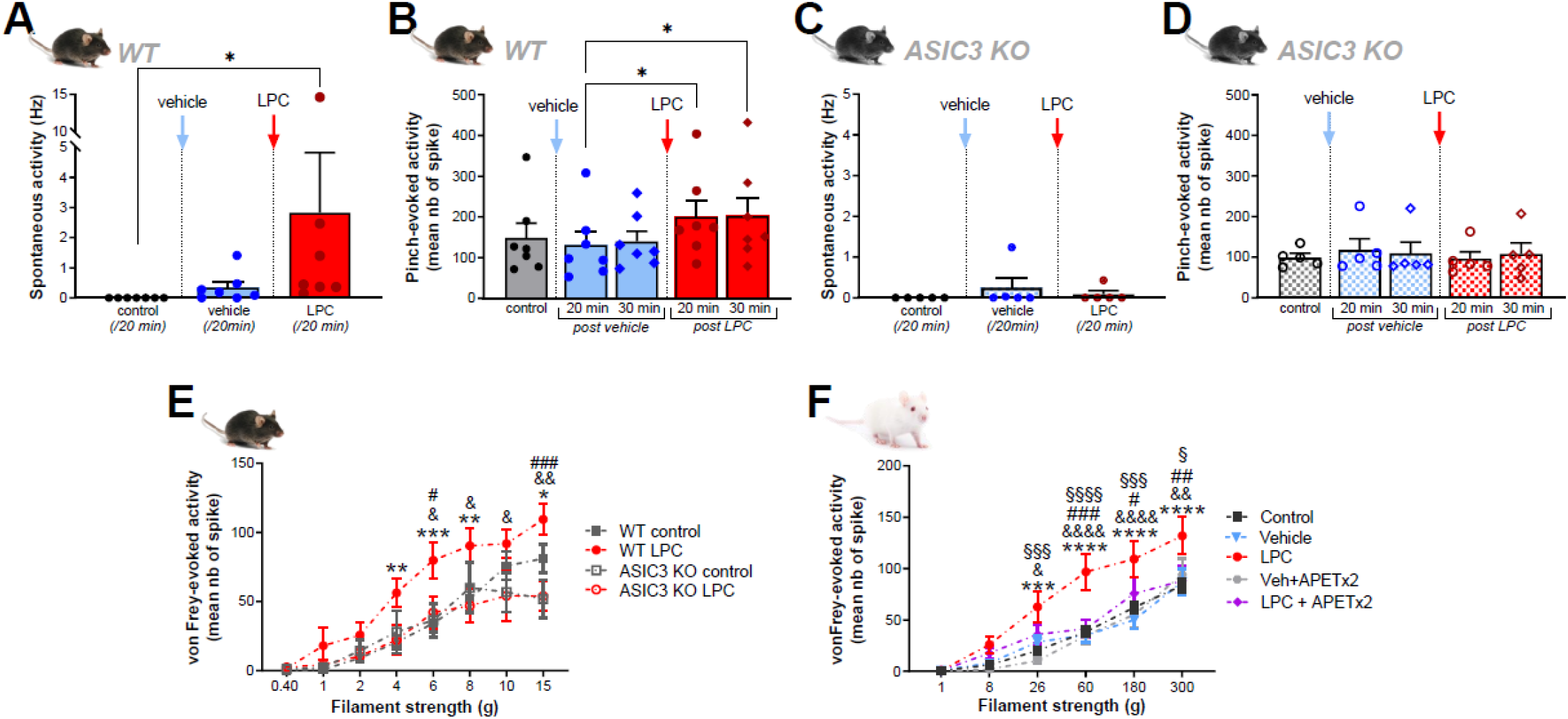
Effect of cutaneous LPC injection on HT neuron activity. **(A)** Global spontaneous discharge of HT neurons before and after vehicle (blue bar and points) and LPC (red bar and point) subcutaneous injection in wild-type mice receptive fields (n=7 neurons from 6 WT mice, Friedman test with p<0.0001 followed by a Dunn’s multiple comparison test: *, p<0.05 for control vs LPC comparison). **(B)** Evoked response of HT neurons to nociceptive stimulation (pinching) in WT mice after vehicle (blue bar and points) and LPC16:0 (red bar and points) subcutaneous injection. Effects were measured 20 min (circle point) and 30 min (diamond point) after respective injections (n= 7 neurons from 6 WT mice, Friedman test with p=0.0025 followed by a Dunn’s multiple comparison test: *, p<0.05). **(C)** Global spontaneous discharge of ASIC3 KO HT neurons before and after vehicle (blue bar and points) and LPC (red bar and points) subcutaneous injection (n=5 neurons from 4 ASIC3 KO mice, no significant difference between vehicle and LPC, p=0.3333, Friedman test). **(D)** ASIC3 KO HT neuron responses to nociceptive stimulation (pinching) before and after vehicle and LPC16:0 subcutaneous injections (n=5 neurons from 4 ASIC3 KO mice, no significant difference, Friedman test with p=0.3848). **(E)** Curves representing the mechanical sensitivity of mouse HT neurons to von Frey filament applications on their receptive fields before (black curves) and after LPC16:0 injection (red curves). Experiments were performed in both WT (full symbols, n=15 neurons in 10 mice and 10 neurons in 6 mice for control and LPC, respectively) and ASIC3 KO (empty symbols, n= 5 neurons) mice (two way ANOVA with p=0.0106 and p<0.0001 for treatment and von Frey filaments effects, respectively, followed by Dunnet’s multiple comparison test: *, p<0.05, **, p<0.01 and ***, p<0.001 for WT LPC vs. WT control; &, p<0.05 and &&, p<0.001 for WT LPC vs. ASIC3 KO LPC; #, p<0.05 and ###, p<0.001 for WT LPC vs. ACIC3 KO control). **(F)** Mechanical sensitivity of rat HT neurons recorded using von Frey filaments before (control, n=39 neurons) and after injection of vehicle (n=10 neurons in 9 rats), vehicle + APETx2 (n=9 neurons in 5 rats), LPC (n=10 neurons in 9 rats) and LPC + APETx2 (n=10 neurons in 6 rats; two way ANOVA with p=0.0004 and p<0.0001 for treatment and von Frey filaments effects, respectively, followed by Dunnet’s multiple comparison test: ***, p<0.001 and ****, p<0.0001 for LPC vs. control; &, p<0.05, &&, p<0.01 and &&&&, p<0.0001 for LPC vs. vehicle; #, p<0.05, ##, p<0.01 and ###, p<0.001 for LPC vs. LPC + APETx2; §, p<0.05, §§§, p<0.001 and §§§§, p<0.0001 for LPC vs. vehicle + APETx2).

The mechanical sensitivity of spinal HT neurons was next assessed using von Frey filaments in mice (Fig. 3E) and rats (Fig. 3F). A set of filaments ranging from 0.4 to 15g was applied consecutively (see Methods) in both WT and ASIC3 KO mice, before and after LPC cutaneous injection into HT neuron receptive fields (Fig. 3E). Before LPC injection (control), HT neurons of both mice genotypes responded roughly in the same way to von Frey stimulations, with classical increase of emitted spikes as a function of filament strength (from 0.31±0.21 to 81.04±10.19 spikes for WT mice, and from 1.2±1.12 to 51.87±13.74 spikes for ASIC3 KO mice), and showed no significant difference in their basal mechanical sensitivities. Following LPC injection, the mechanical sensitivity of WT HT neurons was significantly increased for filaments ≥4g, an effect that was not observed in ASIC3 KO mice (Fig. 3E). The basal von Frey sensitivity of HT neurons in rats, determined with filaments ranging from 1g to 300g, was also enhanced after LPC cutaneous injection (Fig. 3F, from 0.62±0.21 to 81.43±6.38 spikes in control condition, and from 0.87±0.40 to 131.83±18.23 spikes after LPC injection), showing a significant LPC-induced mechanical hypersensitivity for filament strength ≥26g. Similar results were also obtained for rat WDR neurons (Supplementary Fig. 3F). Mechanical hypersensitivity in rats was prevented when the ASIC3 blocker APETx2 was co-injected together with LPC in the receptive fields of both HT neurons (Fig. 3F) and WDR neurons (Supplementary Fig. 3F), further supporting the role of peripheral ASIC3 channels in mediating cutaneous LPC effects.

Altogether, these results demonstrated that a single LPC cutaneous injection was able to induce a short-term (≈45 min) mechanical hypersensitivity of spinal nociceptive neurons in both rats and mice. This effect was observed for painful mechanical stimulations, and was abolished by both genetic (ASIC3 KO) and pharmacological (APETx2) inhibition of ASIC3, strongly supporting the involvement of this channel in LPC-induced mechanical sensitization of spinal nociceptive neurons.

### The LPC-induced sensitization of spinal nociceptive neurons is not modality dependent

Thermal sensitivity of spinal HT neurons was also tested to determine whether sensitization was specific of the stimulus modality. Heat temperature ramps were applied onto rat HT neuron receptive fields, before and after LPC or vehicle injection (Fig. 4A). Cumulative spike number quantification revealed that basal HT neuron activity (control) increased, as expected, as a function of temperature (from 0 spikes at 30°C to 189.20±23.85 spikes at temperatures above 46°C; Fig. 4B). This discharge pattern was significantly enhanced by LPC peripheral injection for temperatures above 42°C, as shown by the increased number of evoked spikes during heat ramp (from 88.75±43.82 spikes at 42°C to 513.05±106.69 spikes at temperatures above 46°C; Fig. 4B), as compared to control (from 2.34±1.35 spikes at 42°C to 189.20±23.85 spikes at temperatures above 46°C; Fig. 4B) and vehicle injection (from 1.06±1.06 spikes at 42°C to 187.61±36.24 spikes at temperatures above 46°C; Fig. 4B). Co-injection of APETx2 prevented LPC-induced thermal hypersensitivity, similarly to what was observed for mechanical hypersensitivity. The thermal-evoked activity of HT neurons was significantly reduced in the LPC+APETx2 condition (from 4.06±3.63 spikes at 42°C to 196.44±37.84 spikes at temperatures above 46°C; Fig. 4B) as compared to LPC alone (from 88.75±43.82 spikes at 42°C to 513.05±106.69 spikes at temperatures above 46°C; Fig. 4B). Finally, a significant decrease of the temperature threshold that triggered HT neuron’s spiking was also observed following LPC cutaneous injection (40.7±0.4°C for LPC *vs*. 44.3±0.2°C and 43.4±0.3°C for control and vehicle, respectively; Fig. 4C), which was abolished by co-injection with APETx2 (44.46±0.4°C for LPC+APETx2; Fig. 4C).

**Figure 4:**
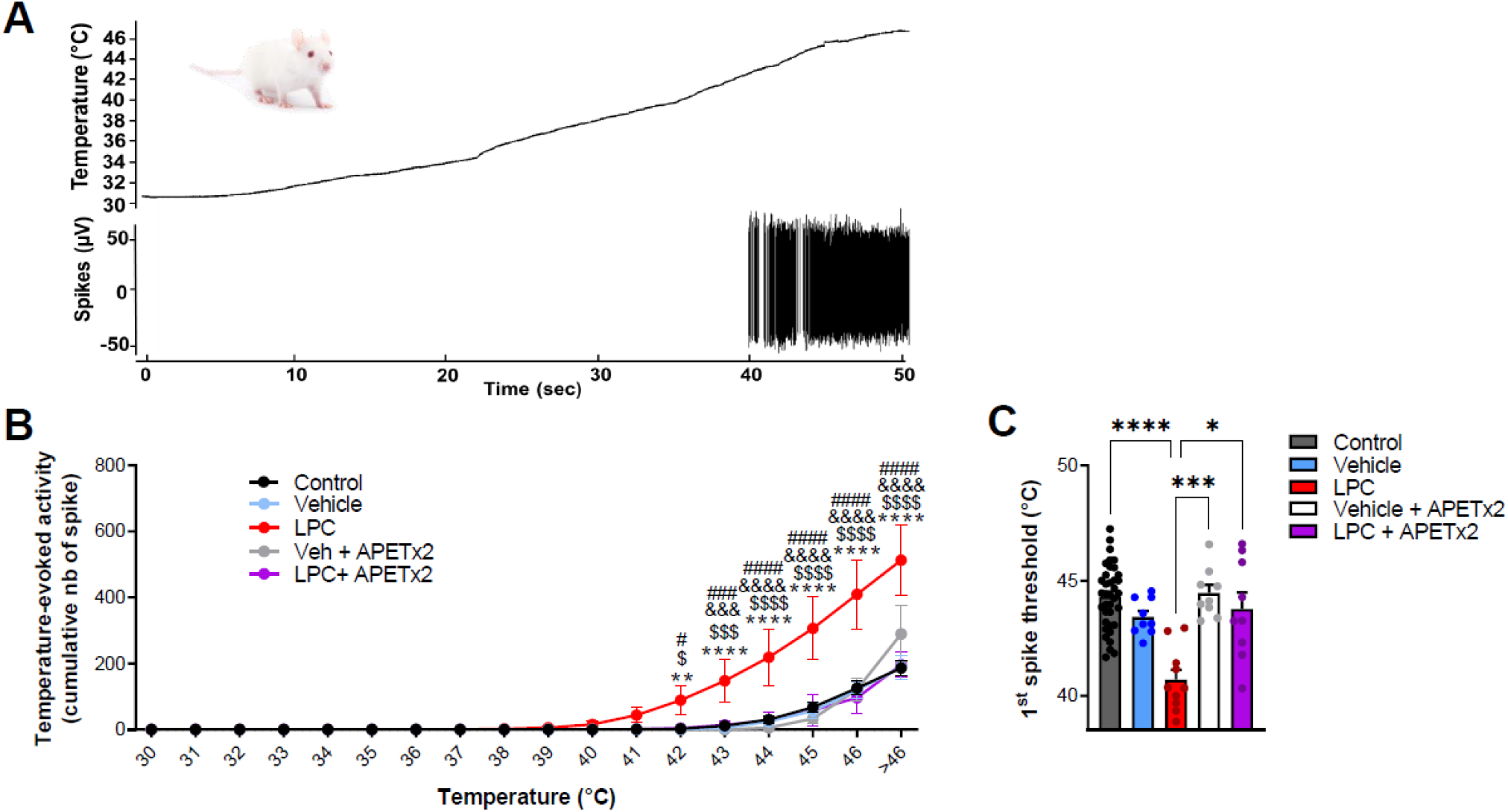
Effect of cutaneous LPC injection on HT neuron thermal sensitivity to heat. **(A)** Example trace of a HT neuron response following heat ramp stimulation. Heat ramps from 30 to 46 degrees were applied onto neuron receptive field (top panel, black curve) while neuron evoked activity was recorded (bottom panel). **(B)** Cumulative representation of the number of spike evoked by heat ramps as a function of the temperature. Experiments were performed before (control, black dots, n=37 neurons from 17 rats) and 20min after cutaneous injection of vehicle (light blue dots, n= 9 neurons from 6 rats), LPC16:0 (red bar dots, n= 10 neurons from 6 rats), vehicle + APETx2 (gray dots, n= 9 neurons from 6 rats) or LPC16:0 + APETx2 (purple dots, n= 9 neurons from 7 rats; two-way ANOVA with p<0.0001 for both treatment and temperature effects, followed by a Dunnet’s multiple comparison test: **, p<0.01 and ****, p<0.0001 for LPC vs. control; $, p<0.05, $$$, p<0.001 and $$$$, p<0.0001 for LPC vs. vehicle; &&&, p<0.001 and &&&&, p<0.0001 for LPC vs. LPC + APETx2; #, p<0.05, ###, p<0.001 and ####, p<0.0001 for LPC vs. vehicle + APETx2). **(C)** Statistical analysis of temperature thresholds that triggered the first spiking activity of HT neurons in response to heat (n=37, 9, 10, 9, 9 and for control, vehicle, LPC, LPC + APETx2 and vehicle + APETx2, respectively, Kruskal-Wallis test with p<0.0001, followed by a Dunn’s multiple comparison test: *, p<0.05, p<0.001 and p<0.0001).

These results showed that LPC also enhanced thermal sensitivity of spinal cord HT neurons, indicating that the LPC-induced short-term sensitization effect was not dependent on the stimulus modality. This effect involved peripheral ASIC3, as for mechanical hypersensitivity.

### Cutaneous LPC injection does not induce enlargement of spinal HT neuron receptive field

To further characterize the mechanism by which peripheral cutaneous LPC affected spinal neurons and to determine whether it could induce some spinal sensitization features, experiments were performed to study HT neuron receptive field areas in rats, *i*.*e*., to see how they could possibly be affected by LPC (Fig. 5A). Enlargement of the receptive field of spinal cord neurons is typically associated with central sensitization (Latremoliere and Woolf, 2009).

**Figure 5:**
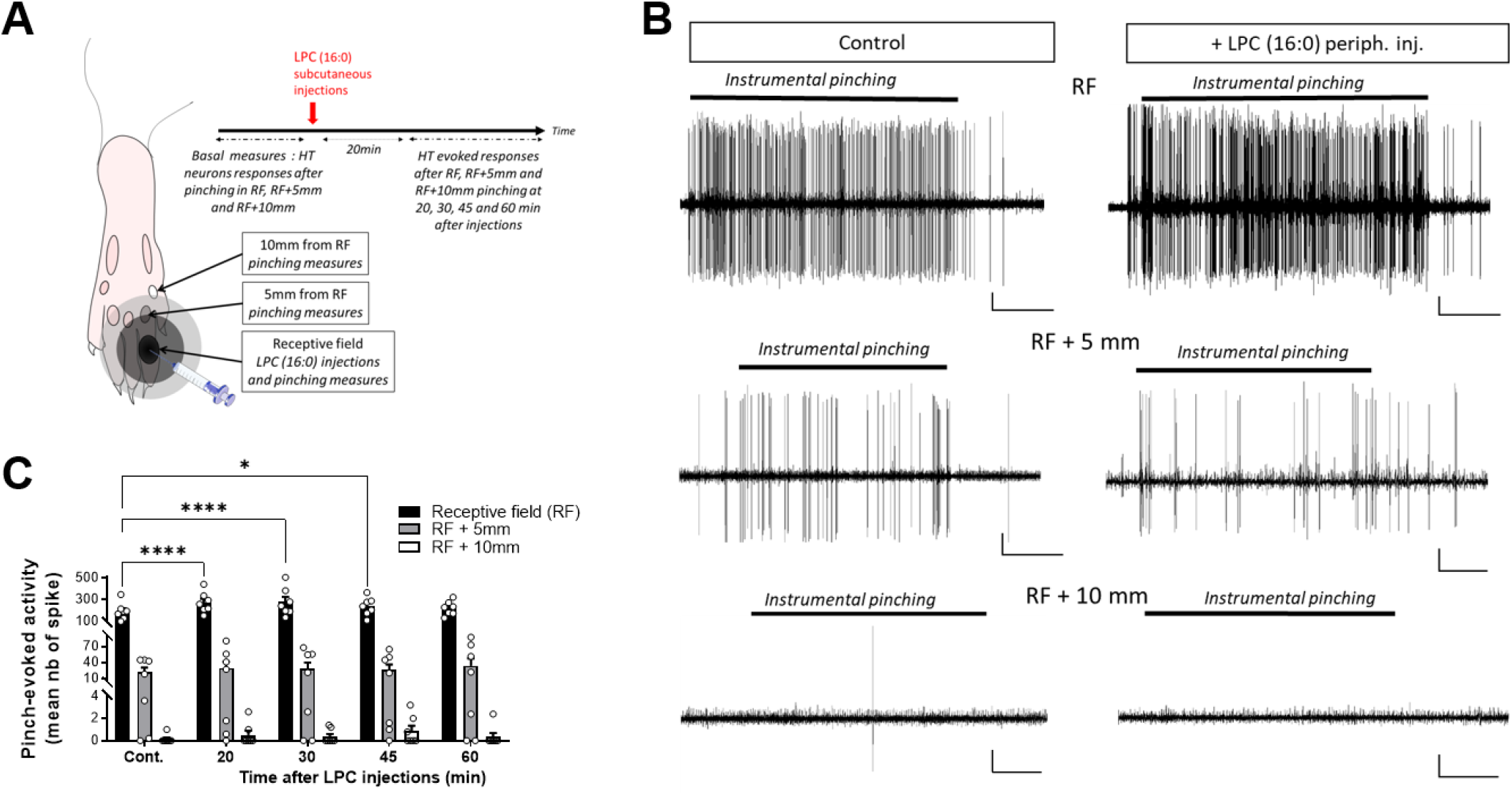
Effect of cutaneous LPC injection on HT neuron receptive field. **(A)** Experimental protocol used to test the possible enlargement of HT neuron receptive field (RF). Different areas were initially determined: RF, RF+5mm and RF+10mm. Noxious pinches (instrumental pinching, 300g) were then applied onto these different areas. **(B)** Representative recordings of a HT neuron discharge following pinching of RF, RF+5 mm or RF+10mm areas before (left panels) and 20min after LPC16:0 peripheral injection (right panels, scale: 10µV – 1 sec). **(C)** HT neuron responses following RF (black bar), RF+5mm (grey bar) and RF+10mm (white bar) noxious stimulations, before and after LPC16:0 cutaneous injection. LPC only enhanced HT neurons evoked response when pinches were applied within RF and not at RF+5mm or RF+10mm (n=7 neurons from 7 rats, two-way ANOVA with p=0.0043 and p<0.0001 for time of LPC and RF area effects, followed by a Dunnet’s multiple comparison test: *, p<0.05 and ****, p<0.0001 as compared to control).

Receptive field areas were first determined before LPC injection as the areas inside which noxious pinching induced strong responses of HT neurons (Fig. 5A-B, upper-left trace). Noxious stimulations were then applied 5mm and 10mm outside the receptive field areas (Fig. 5A). HT neuron responses to pinch were weak 5mm and absent 10mm away from the receptive field (Fig. 5B, middle and bottom left traces). Following LPC cutaneous injection, HT neuron responses were significantly increased when receptive field was stimulated (Fig. 5B, upper-right trace and Fig. 5C), as observed previously (Figs 1 and 3), but no significant change was observed when stimulations were applied 5mm or 10mm away from receptive field (Fig. 5B, middle- and bottom-right traces, and Fig. 5C). Accordingly, cutaneous LPC injection did not induce enlargement of spinal HT neuron receptive field, which is not consistent with development of central sensitization.

## DISCUSSION

We already demonstrated that cutaneous LPC, co-injected together with AA, produced acute pain behavior in rodent (Marra *et al*., 2016). The aim of this work was to *(i)* study the pain message generated by LPC when injected alone in the skin, *(ii)* determine the contribution of peripheral ASIC3 to the generation of this message and, *(iii)* investigate how this message is processed by dorsal spinal cord neurons.

Our data show that cutaneous LPC injection exclusively affects the nociceptive pathway, inducing an ASIC3-dependent sensitization of peripheral C-fibers associated with increased spontaneous and evoked-activities of spinal HT and WDR neurons. The activity of spinal LT neurons remained unaffected by cutaneous injection of LPC, consistent with a lack of effect of the lysolipid on non-nociceptive Aβ fibers. The sensitizing effect of LPC on HT and WDR neurons occurs following a single cutaneous injection, it lasts approximately 45 minutes and it is modality-independent since neuronal responses to noxious heat and mechanical stimulations are both potentiated. Basal mechanical and thermal sensitivity is not affected in our experimental conditions by pharmacological or genetic inhibition of ASIC3 with APETx2 and knockout, respectively, in agreement with previous data using ASIC3 KO mice to test behavioral responses following noxious thermal or von Frey mechanical stimulations (Borzan *et al*., 2010; Chen *et al*., 2002; Price *et al*., 2001). On the other hand, the increased C-fiber mechanical and thermal sensitivity induced by peripheral injection of LPC is clearly dependent on the ASIC3 activity. It is also fully consistent with the recent description of mechanical hypersensitivity after cutaneous injection of LPC in mice (Rimola *et al*., 2020). The persistent, depolarizing ASIC3 current generated by LPC is likely to participate in this ASIC3-dependent sensitization of C-fibers, as demonstrated in primary cultures of dorsal root ganglia neurons (Deval *et al*., 2003). The augmented peripheral nociceptive inputs drive the increase in spinal outputs from HT and WDR neurons.

An increased spinal output can result from both peripheral and central sensitization processes. Spinal cord neurons subject to central sensitization exhibit typical features, such as an increase of spontaneous activity, a reduction in the threshold for activation by peripheral stimuli, an increased response to suprathreshold stimulations, and an enlargement of the neuron receptive field (Latremoliere and Woolf, 2009). We demonstrate here that HT and WDR spinal neurons exhibited enhanced spontaneous activities following LPC cutaneous injections, and LPC also reduced the temperature threshold that triggered spiking activity in HT neurons. Moreover, the facilitation process of windup was also potentiated by LPC. On the other hand, experiments using von Frey filaments did not reveal any clear significant appearance of HT and WDR neurons responses to subthreshold stimulations. Finally, the increased response of spinal neurons to suprathreshold stimuli is rather short (45 minutes) and, importantly, there was no enlargement of spinal neuron receptive fields. The effects of cutaneous LPC on spinal neuron activities, *i*.*e*., increased of both spontaneous firing and evoked responses to noxious stimuli, seem therefore not to involve central sensitization process, but rather an ASIC3-dependent short-term sensitization of peripheral nociceptive inputs.

LPC displays a good specificity for ASIC3 compared to other pain-related ASICs also expressed in peripheral sensory neurons such as ASIC1a and ASIC1b, as shown here and in Marra et al (2016). However, LPC not only modulates ASIC3 (Marra *et al*., 2016), but has also been reported to affect other pain-related channels including TREK1 (Maingret *et al*., 2000), TRPM8 (Andersson *et al*., 2007; Gentry *et al*., 2010), TRPC5 (Flemming *et al*., 2006; Sadler *et al*., 2020), and more recently TRPV1 (Rimola *et al*., 2020). Our data demonstrate an overall excitatory effect of peripheral injection of the lysolipid on spinal neurons and confirm the excitatory nature of LPC. They also support an important role for ASIC3 in this effect at least in naïve animals, which does not preclude the involvement of other LPC-modulated channels to its cutaneous effects *in vivo* in physiological or pathophysiological conditions.

This study demonstrates how a single cutaneous injection of LPC can generate a short-term sensitization of peripheral nociceptive fibers. The underlying mechanism involves pain-related ASIC3 channels, which can be constitutively activated by this lipid. The nociceptive input induced by a single LPC cutaneous injection is clearly not sufficient to drive a spinal synaptic plasticity strong enough to lead to central sensitization. However, ASIC3 has been identified as a triggering factor of chronic muscle and joint pain in animal models (Hung *et al*., 2020; Sluka *et al*., 2003; Sugimura *et al*., 2015), and the high level of LPC detected in patients with joint or muscle painful diseases (Hung *et al*., 2020; Marra *et al*., 2016) could definitely support the development of central sensitization underlying chronic pain through the same mechanisms as those described here.

## METHODS

### Cell culture and transfections

HEK293 cell line was grown in DMEM medium supplemented with 10% of heat-inactivated fetal bovine serum (BioWest) and 1% of antibiotics (penicillin + streptomycin, BioWhittaker). One day after plating, cells were transfected with either pIRES2-rASIC1a-EGFP (rat ASIC1a), pIRES2-rASIC1b-EGFP (rat ASIC1b) or pIRES2-rASIC3-EGFP (rat ASIC3) vectors using the JetPEI reagent according to the supplier’s protocol (Polyplus transfection SA, Illkirch, France). Fluorescent cells were used for patch-clamp recordings 2–4 days after transfection.

### Patch clamp experiments

Whole cell configuration of the patch clamp technique was used to record membrane currents at a holding potential of −80mV (voltage clamp mode). Recordings were made at room temperature using an axopatch 200B amplifier (Axon Instruments) with a 2 kHz low-pass filter. Data were digitized by a Digidata 1550 A-D/D-A converter (Axon Instruments), sampled at 20 kHz and recorded on a hard disk using pClamp software (version 11; Axon Instruments). The patch pipettes (2–6 MΩ) were filled with an intracellular solution containing (in mM): 135 KCl, 2 MgCl_2_, 5 EGTA, and 10 HEPES (pH 7.25 with KOH). The extracellular solution bathing the cells contained (in mM) the following: 145 NaCl, 5 KCl, 2 MgCl_2_, 2 CaCl_2_, 10 HEPES (pH 7.4 with N-methyl-D-glucamine). ASIC currents were induced by shifting one out of eight outlets of a homemade microperfusion system driven by solenoid valves, from a holding control solution (*i*.*e*., pH 7.4) to an acidic test solution (pH 7.0 or pH 6.6). Cells were considered as positively transfected when they exhibited a visible GFP fluorescence and a significant pH6.6-evoked current of at least 300pA (IpH6.6 ≥ 300pA). Non-transfected (NT) cells were used as controls and they were selected in Petri dishes having undergone the transfection protocol described above, but with no visible GFP fluorescence and no significant pH-6.6-evoked current.

### Animals

Experiments were performed on adult male Wistar Han rats (Charles River, age > 6 weeks), on adult male C57Black6J wild-type mice (WT, Janvier Lab, age > 7 weeks) and on ASIC3 knockout mice (ASIC3 KO, internal animal husbandry, age > 7 weeks). The protocol was approved by the local ethical committee and the French government (agreement n° 02595.02). Animals were kept with a 12-h light/dark cycle with *ad libitum* access to food and water and were acclimated to housing and husbandry conditions for at least 1 week before experiments.

### Surgery

Anesthesia was induced in a box ventilated with a mix of air and isoflurane 4% (Anesteo, France). Animals were then placed on a stereotaxic frame (M2E, France) and were kept under anesthesia using a mask diffusing a mix of oxygen and isoflurane 2%. The head and vertebral column of the animal under experimentation was stabilized by ear bars and vertebral clamps, respectively, while a limited laminectomy was perform between vertebrae T13 and L2. Dura was carefully removed, and the spinal cord was immerged with artificial cerebrospinal fluid (ACSF containing 119 mM NaCl, 2.5mM KCl, 1.25 mM NaH_2_PO_4_, 1.3 mM MgSO_4_, 2.5mM CaCl_2_, 26 mM NaHCO_3_, 11 mM glucose and 10 mM HEPES, pH adjusted to 7.4 with NaOH) before starting electrophysiological recordings.

### In vivo electrophysiological recordings of spinal cord neurons

Single-unit extracellular recordings of spinal cord dorsal horn neurons were made with tungsten paralyn-coated electrodes (0.5 MΩ, WPI, Europe) and using Spike2 acquisition system (Cambridge Electronic Design, UK). The tip of a recording electrode was initially placed at the dorsal surface of spinal cord using a micromanipulator (M2E, France) and this initial position determined the zero on the micromanipulator’s micrometer. The electrode was then progressively moved down into the dorsal horn until the receptive field of a spinal neuron was localized on the ipsilateral plantar hindpaw using mechanical stimulations. Neuronal signals were bandpass filtered (0.3-30 kHz) and amplified using a DAM80 amplifier (WPI, Europe), digitized with a 1401 data acquisition system (Cambridge Electronic Design, Cambridge, UK), sampled at 20 kHz and finally stored on a computer.

Once a spinal neuron was isolated with its receptive field, non-noxious (brushing) and noxious (pinching) stimulations were used to characterize the neuron type (Figs 1B-C and 2A). Classically, spinal neurons were differentiated depending on the peripheral input they receive: *(i)* LT neurons (Fig. 1B), receiving input from Aβ-peripheral fibers and strongly responding to non-noxious brushing (Fig. 1B1), but not to noxious pinching (Fig. 1B2), *(ii)* HT neurons (Fig. 1C), receiving input from Aδ/C-fibers and only responding to noxious pinching by a sustained activity that lasts the whole time of stimulation (Fig. 1C1 & C2), and *(iii)* WDR neurons (Fig. 2A), responding to both noxious and non-noxious stimulations (Aβ, Aδ and C inputs, Fig. 2A1 & A2), and known to exhibit a facilitation process (Latremoliere and Woolf, 2009) called windup (Fig. 2A3 & A4). The depths of neurons recorded in this study ranged from 0-250µm, 200-500µm and 350-1100µm for LT, HT and WDR neurons, respectively.

### Stimulation protocols of spinal neuron receptive fields

Receptive fields of spinal neurons were stimulated every 10 minutes by applying 10 consecutive brushings, using paint brush n°10, and/or 5 consecutive pinches, using either calibrated forceps (300g stimulations, Bioseb, France) or classical forceps (Moria MC40/B, Fine Science Tools, Germany) for rats and mice, respectively. These stimulation protocols were used to generate Aβ-evoked response to non-noxious brushings and Aδ/C-evoked responses to noxious pinchings. In addition to these mechanical stimuli, receptive fields of WDR neuron also received repetitive electrical stimulations (protocol of 16 supraliminar 4ms pulses, Dagan S900 stimulator) to induce windup. Intensity of currents injected for windup was determined as the intensity required to evoke less than 10 action potentials (APs) at the first stimulation, corresponding to 1.2-3 times the AP thresholds.

The evoked response of spinal neurons to mechanical non-noxious and noxious stimulations was also measured using von Frey filaments. Different filaments were used to determine the mechanical sensitivity of HT and WDR neurons in rats (1g ; 8g ; 26g ; 60g ; 180g ; 300g) and in mice (0.40g, 1g, 2g, 4g, 6g, 8g, 10g, 15g). Each filament was applied three times during three seconds.

Finally, the response of spinal neurons to thermal stimulation was studied by applying heat ramps onto animal hindpaws. Heat ramps were applied by running a trickle of warm water on the neuron’s receptive fields using a temperature controller (CL-100, Warner instruments, USA). A temperature probe was placed on the center of the receptive fields and temperature was monitored for the entire experiment duration. Temperature was initially set at 30°C and heat ramps were delivered for 47 seconds up to 47°C, every 10 minutes, before and after injection of LPC or vehicle.

### Peripheral injection of lipids and drugs

Subcutaneous injection was done in the receptive fields of spinal neurons (20µl and 10µl for rats and mice, respectively) using a 28 gauge needle connected to a 50µl Hamilton syringe. LPC16:0 was purchased from Anvanti (Coger, France), prepared as stock solutions in ethanol, and injected either alone (4.8 nmole and 9.6 nmole diluted NaCl 0.9% for rats and mice, respectively) or in combination with APETx2 (0.2nmoles ; purchased from Smartox Biotechnology, France, and prepared as stock solution in NaCl 0.9%). Ethanol 0.24 or 0.96 % were used as vehicle solutions for rats and mice, respectively.

### Spike sorting and analysis

Off-line analyses of *in vivo* electrophysiological recordings were done using Spike2 (Cambridge Electronic Design, UK) and Matlab (MathWorks, Natick, MA, USA) softwares. The spike sorting was first performed with Spike2, using principal component analysis of spike waveforms. The spikes and associated stimulation train were then exported to Matlab to perform further analysis. For each neuron, both spontaneous activities and evoked responses to noxious or non-noxious stimulations were quantified as the number of spike emitted at rest and during the different stimulations, respectively. Matlab codes were used to calculate mean number of spikes. Spontaneous activities were calculated over 20 min periods starting immediately after vehicle, LPC, vehicle + APETx2 or LPC + APETx2 injections. For non-noxious brushings, the mean number of spikes was calculated over 10 consecutive stimulations. For noxious pinching, the mean number of spikes was calculated over the 5 consecutive 5-sec stimulations. For windup analysis of WDR neurons, each interval between repetitive electrical stimulations was divided into periods, so that the spikes evoked by Aδ and C-fibers can be distinguished (Fig. 2 and Supplementary Fig. 2A). Indeed, spikes emitted within the 20-90 ms interval after the stimulation artefact were attributed to Aδ-fibers, whereas those emitted during the 90-1,000ms interval were attributed to C-fibers (90-350 ms) and the after depolarization (AD) period (350-1,000 ms). Windup curves were established by counting the number of spikes emitted during C-fiber + AD periods for each of the 16 repetitive electrical stimulations. Windup was then expressed as the area under curve (AUC) for each curve. AUC was calculated from the baseline corresponding to the first number of spikes.

For experiments using von Frey filaments, the mean of emitted spikes number was calculated over three consecutive 3-sec stimulations.

### Statistical analysis of data

Graphs and statistical analysis were done using GraphPad Prism software (GraphPad Software, San Diego, CA). Numerical values are given as mean ± SEM, unless otherwise stated. Statistical differences between sets of data were assessed using either parametric or nonparametric tests followed by *posthoc* tests, when appropriate. Statistical test used and significant *p*-values are indicated in each figure legend.

## ACKNOWLEDGEMENTS

We thank Drs A. Baron, S. Diochot, J. Noël, M. Salinas and P. Inquimbert for helpful discussions, V. Friend and J. Salvi-Leyral for technical support, and V. Berthieux for secretarial assistance. This work was supported by the Centre National de la Recherche Scientifique, the Institut National de la Santé et de la Recherche Médicale, the Association Française contre les Myopathies (AFM grant #19618), the Agence Nationale de la Recherche (ANR-11-LABX-0015-01 and ANR-17-CE16-0018) and the NeuroMod Institute of University Côte d’Azur (UCA).

## AUTHOR CONTRIBUTIONS

LP, Conception and design, acquisition, analysis and interpretation of in vivo data. KD, Acquisition, analysis and interpretation of in vitro data. EL, Interpretation of data. ED, Conception and design, analysis and interpretation of data. All the authors have contributed to drafting the manuscript and have given approval to its deposition on BiorXiv.

