## supplementary figures for "ASIC3-dependent spinal cord nociceptive signaling in cutaneous pain induced by lysophosphatidyl-choline"

### SUPPLEMENTARY MATERIAL

***In vitro effects of LPC on ASICs***

LPC was superfused for 30 seconds onto patch-clamped cells, while fast extracellular pH drops from pH7.4 to pH7.0 were made to generate acid-evoked currents every ten seconds (Supplementary Fig. 1A) to record both the activating and potentiating effects of LPC on ASIC3 (1 and 2 on Supplementary Fig. 1A; (Marra *et al.*, 2016). At resting physiological pH7.4, LPC also dose- and time-dependently activated a small but significant non-inactivating current in ASIC3-transfected cells (Supplementary Fig. 1A and 1C), as compared to non-transfected cells (NT). This non-inactivating ASIC3 current started to activate after 10- to 20-seconds of LPC superfusion and reached its higher amplitude after 30 seconds (Supplementary Fig. 1C). A significant potentiation of the pH7.0-evoked ASIC3 current was observed for LPC concentrations of 5µM and 10µM, starting after 10 seconds of superfusion and reaching the strongest effect after 30 seconds (Supplementary Fig. 1D).

LPC was also tested on ASIC1a and ASIC1b channels, which are also highly expressed in sensory neurons (Chen *et al.*, 1998; Waldmann *et al.*, 1997a; Waldmann *et al.*, 1997b). Due to the lower pH sensitivities and longer reactivation kinetics of ASIC1a and ASIC1b, the protocol was adapted so that the pH drops were made from pH7.4 to pH6.6 every 60 seconds (Supplementary Fig. 1B). No activation or potentiation effect of LPC (5µM) was observed on ASIC1a or ASIC1b, except a very small but statistically significant potentiation of the pH6.6-evoked ASIC1b current (Supplementary Fig. 1C and 1D), demonstrating the specificity of the lysolipid for ASIC3 within the ASIC channel family.

Altogether, these data confirm the activating/potentiating effects of LPC on ASIC3 currents for concentrations in the 1-10µM range.

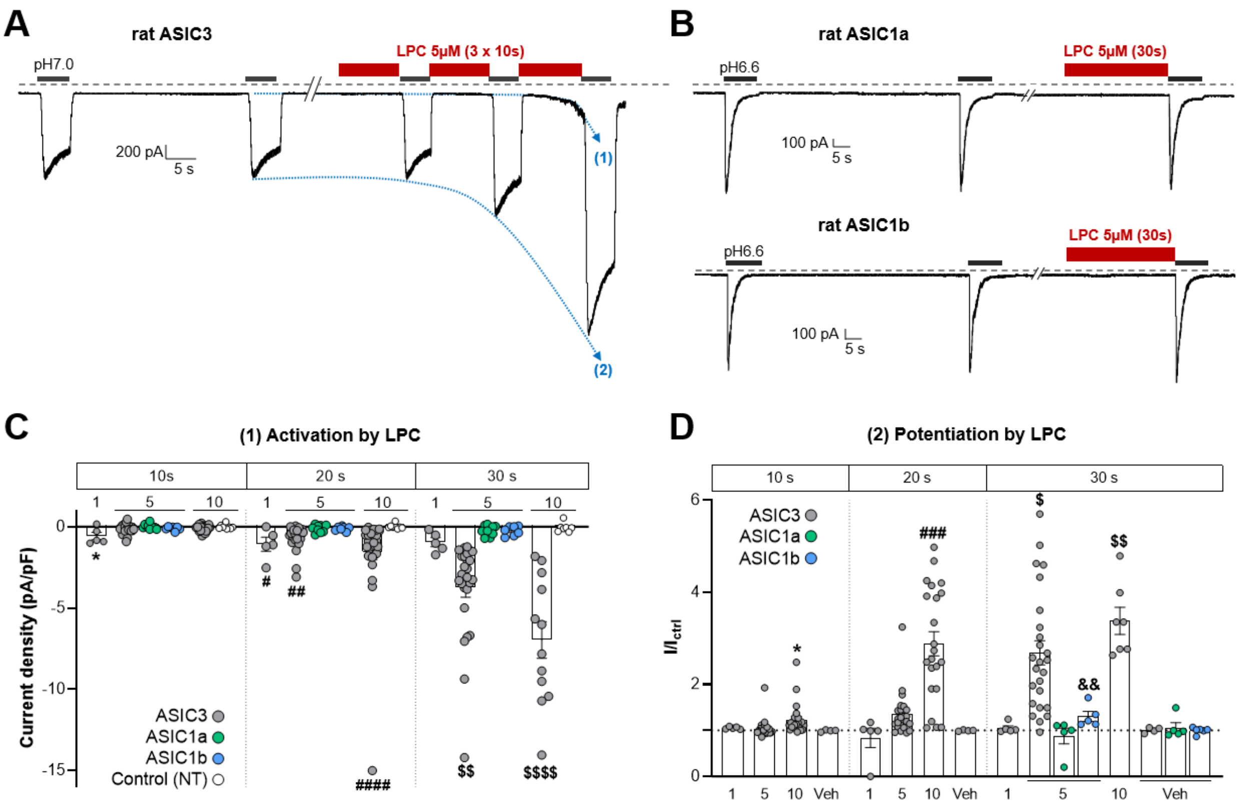


***Supplementary Figure 1*: In vitro effects of LPC on ASIC1 and ASIC3 channels. (A-B)** *Representative current traces recorded at -80mV from HEK293 cells transfected with either rat ASIC3 (A), rat ASIC1a (B, upper trace) or rat ASIC1b (lower trace) subunits. Currents were elicited by repetitive extracellular pH drops (from pH7.4 to pH7.0 every 10 s for ASIC3, and from pH7.4 to pH6.6 every 60 s for ASIC1a and ASIC1b) and LPC16:0 (5µM) was applied extracellularly for 30 seconds, as indicated above each traces. These protocols allowed studying the effects of LPC on both the basal activation (1) and the potentiation (2) of acid-evoked ASIC currents.* ***(C)*** *Bar graphs showing the mean currents activated by different concentrations of LPC16:0 (1µM, 5µM and 10µM) after 10-, 20- and 30-second application on ASIC1a, ASIC1b and ASIC3 transfected cells (n=5-30; *, p<0.05 as compared to control non-transfected cells (NT) at 10s ; #, p<0.05, ## , p<0.01 and ####, p<0.0001 as compared to control NT at 20s ; $$ , p<0.01 and $$$$, p<0.0001 as compared to control NT at 30s, Kruskal-Wallis tests followed by Dunn's multiple comparison tests).* ***(D)*** *Bar graphs showing the potentiation of pH-evoked ASIC currents (pH7.0 for ASIC3 and pH6.6 for ASIC1a and ASIC1b) by different concentrations of LPC16:0 (1µM, 5µM and 10µM) after 10-, 20- and 30-second application on ASIC1a, ASIC1b and ASIC3 transfected cells. The pH-evoked current amplitudes are normalized ($I/I_{ctrl}$) to the current measured before extracellular application of LPC (n=4-25 ; *, p<0.05 as compared to control vehicle at 10s ; ###, p<0.001 as compared to control vehicle at 20s ; $ , p<0.05 and $$, p<0.01 as compared to control vehicle for ASIC3 at 30s; &&, p<0.01 as compared to control vehicle for ASIC1b at 30s, Kruskal-Wallis tests followed by Dunn's multiple comparison tests).*

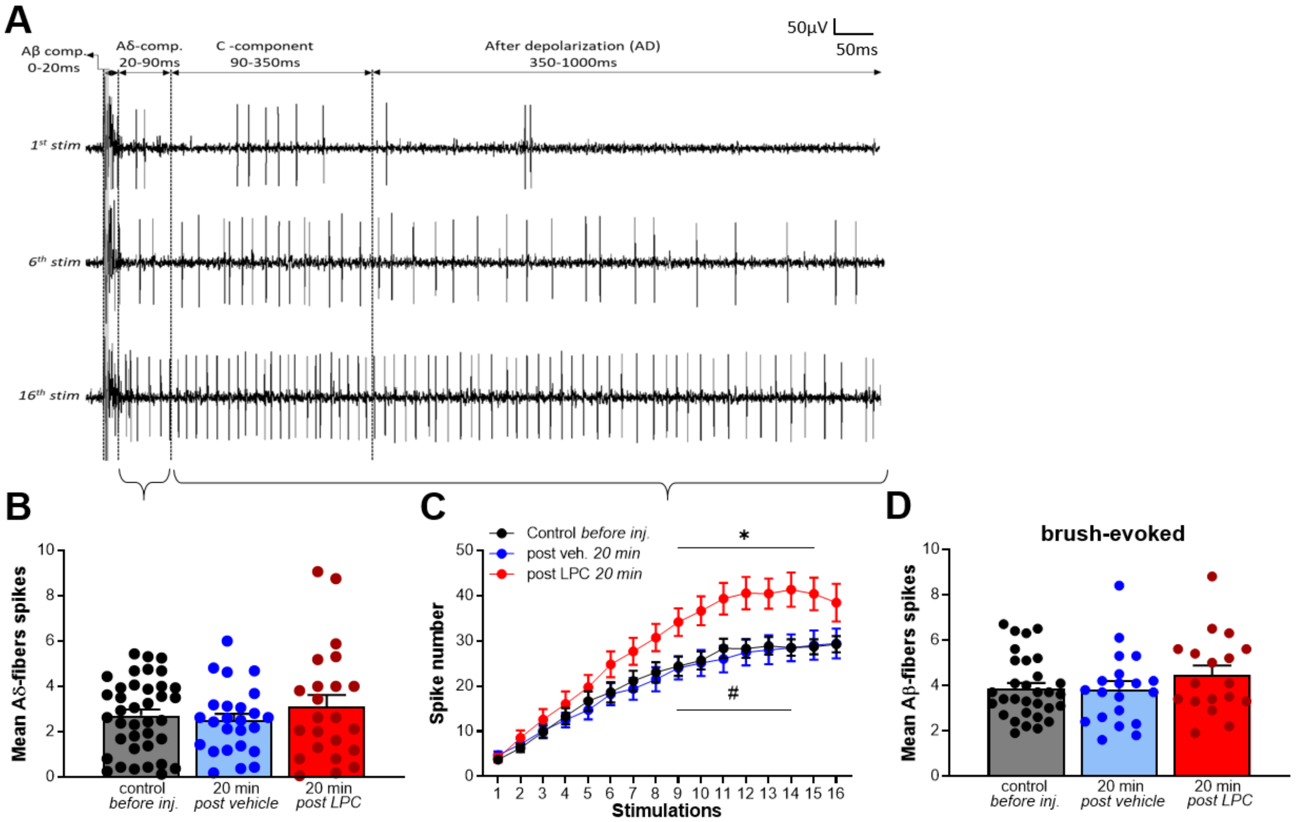


***<u>Supplementary Figure 2</u>: Effect of LPC cutaneous injection on WDR neuron activity evoked by Aβ, Aδ and C-fibers. (A)*** *Typical recordings obtained from a WDR neuron after the 1st (top trace), the 6th (middle trace) and the 16th (bottom trace) electrical stimulation of its receptive field. Windup was induced by 16 supra-threshold electrical stimulations at 1Hz (see Methods). Vertical dotted lines delimit the four periods used to distinguish the WDR spiking evoked by Aβ (0-20ms), Aδ (20-90ms) and C fibers (90-350ms), and with the spikes emitted during the 350-1,000 ms interval attributed to the after depolarization (AD) period. If Aβ-evoked activity is difficult to distinguish because emitted spikes are mixed with stimulation artifacts, activities related to Aδ and C-fibers can easily be measured.* ***(B-D)*** *Pooled data regardless of the protocol used (1 and 2, see fig1A) Part of these data were already presented in Fig. 2.* ***(B)*** *Twenty minutes after its cutaneous injection, LPC had no significant effects on the WDR spiking activities evoked by Aδ inputs (n=38, 25 and 23, for control, vehicle and LPC, respectively; Kruskal-Wallis test with p=0.8026).* ***(C)*** *Windup curves representing the number of emitted spikes during the C-fiber + AD periods as a function of the electrical stimulation number, before and 20 min after vehicle or LPC cutaneous injection (n=38, 25 and 23 for control, vehicle and LPC, respectively; two-way ANOVA test with p=0.0128 and p<0.0001 for the treatment and the stimulation number, respectively; *, p<0.05 and #, p<0.05 for control vs. LPC, and vehicle vs. LPC, respectively).* ***(D)*** *Mean number of spikes evoked by brushing and related to Aβ-fibers (n=31, 20 and 18, for control, vehicle and LPC, respectively; Kruskal-Wallis test with p=0.4528).*

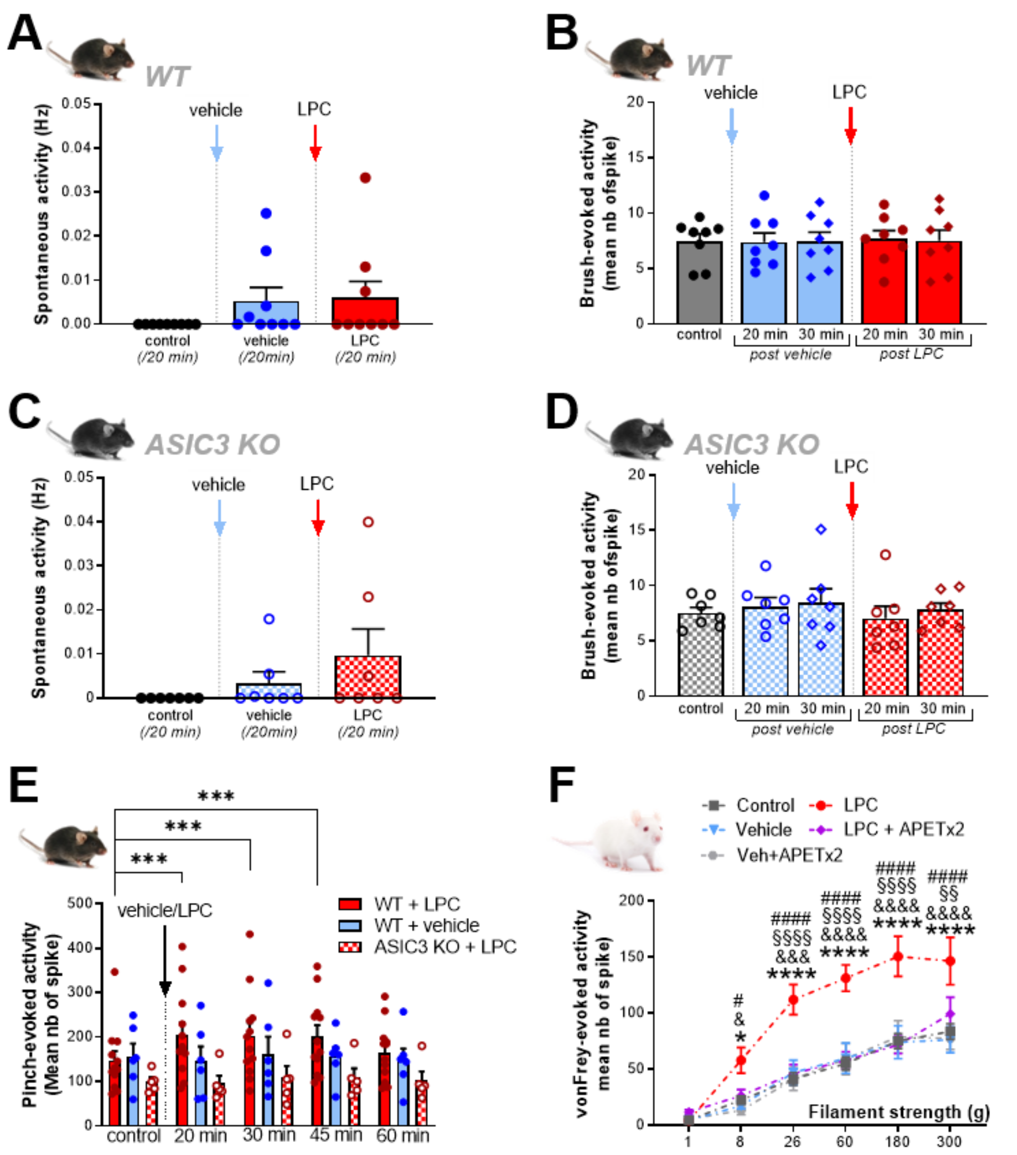


**Supplementary Figure 3:** ***Effect of LPC cutaneous injection on LT, HT and WDR neuron mechanical sensitivity.*** ***(A)*** *Global spontaneous discharge of LT neurons (n= 9 neurons from 5 WT mice) before and after vehicle (blue bar and points) and LPC16 :0 (red bar and points) subcutaneous injection in the hindpaw receptive fields (no-significant difference, Friedman test with p=0.1728).* ***(B)*** *Evoked response of LT neurons to non-nociceptive stimulation (brushing) before and after vehicle (blue bar and points) and LPC16:0 (red bar and points) subcutaneous injection. Effects were measured at 20 min (circle point) and 30 min (diamond point) after injection (n=8 neurons from 4 mice, Friedman test with p=0.8687).* ***(C-D)*** *Spontaneous activity (C) and brush-evoked responses (D) of ASIC3 KO LT neurons (n=7 neurons from 4 mice, no significant difference with p=0.1944 (C) and p=0.1751 (D), Friedman tests).* ***(E)*** *Duration of LPC effect on the brush-evoked responses of mouse HT neurons. Part of the data were pooled from different experiments already presented in Fig. 2, regardless of the protocol used (n=12 neurons from 10 mice, 5 neurons from 4 mice and 6 neurons from 6 mice for WT + LPC, ASIC3 KO + LPC and WT + vehicle, respectively; two-way ANOVA test with p=0.0976 and p=0.1197 for time and LPC treatment effects, respectively, followed by a Dunnet's multiple comparison test: ***, p>0.001).* ***(F)*** *Mechanical sensitivity of rat WDR neurons obtained using von Frey filaments before (control, n=33 neurons from 24 rats) and after injection of vehicle (n=8 neurons from 5 rats), LPC (n=8 neurons from 7 rats), LPC + APETx2 (n=9 neurons from 6 rats) and vehicle + APETx2 (n=8 neurons from 6 rats ; two-way ANOVA with p<0.0001 for both treatment and von Frey filament effects, followed by Dunnet's multiple comparison test: *, p<0.05 and ****, p<0.0001 for LPC vs. control; &, p<0.05, &&&, p<0.001 and &&&&, p<0.0001 for LPC vs. vehicle; §, p<0.05, §§, p<0.01 and §§§§, p<0.0001 for LPC vs. LPC + APETx2; #, p<0.05 and ####, p<0.0001 for LPC vs. vehicle + APETx2).*
